# A truncation library for the soluble production of squalene synthase from *Candida albicans* in *Escherichia coli*

**DOI:** 10.1101/2025.02.06.636915

**Authors:** Carmien Tolmie, Diederik J. Opperman

## Abstract

The rising global burden of infectious fungal disease and the steady increase of resistance by fungal pathogens to available treatments necessitate a concerted effort to develop new antifungal medicines. Squalene synthase is a critical enzyme in the synthesis of ergosterol, a fungal-specific sterol, and has not yet been exploited as an antifungal drug target for pathogenic yeast. In this study, we heterologously produce squalene synthase (SQS) from the opportunistic pathogen *Candida albicans* in *Escherichia coli* to obtain high-quality pure protein for structural studies. A library of nine C- or N&C-terminally truncated SQS variants was successfully produced and purified with immobilised metal-affinity chromatography (IMAC). Two variants were upscaled and purified to near-homogeneity, but size-exclusion chromatography showed possible aggregation. Nevertheless, this study provides a starting point to further optimise buffer conditions to produce high-quality heterologous SQS from *C. albicans* for downstream crystallography experiments.

## Introduction

Squalene synthase (SQS, EC 2.5.1.21) catalyses the first committed step in the biosynthesis of sterols in eukaryotes, and is the branch point from the production of non-steroidal isoprenoid molecules in the mevalonate pathway (Robinson et al., 1993; Thompson et al., 1998). In this reaction, SQS catalyses the head-to-head dimerisation of two molecules of farnesyl pyrophosphate, followed by NADPH-dependent reduction to squalene (Figure 1). Animals produce cholesterol as the main sterol, while fungi, protozoa and plants produce ergosterol. These counterpart sterols are essential in maintaining membrane fluidity for the survival of eukaryotic cells and have various other critical roles, such as lipid metabolism and the production of bile salts (as in the case of cholesterol). Additionally, ergosterol is an immunoactive molecule that plays a role in the pathogenesis of fungi (Rodrigues, 2018).

**Figure 1.**
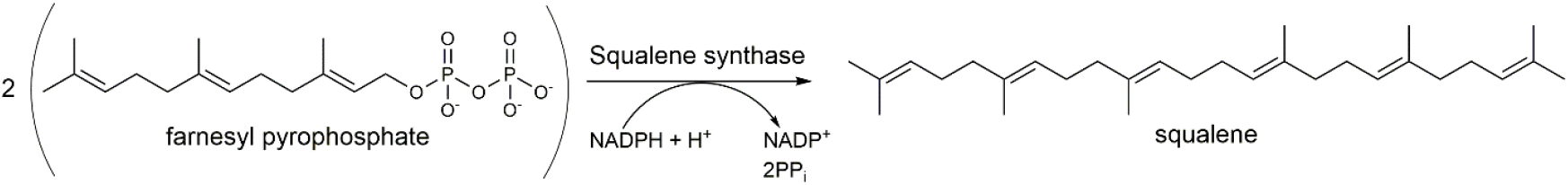
Conversion of two molecules of farnesyl pyrophosphate to squalene by squalene synthase in the sterol biosynthesis pathway.

In humans, SQS has been a target for ongoing research to develop cholesterol-lowering drugs (Haginoya et al., 2024; Liu et al., 2012; Matralis et al., 2023). In *Trypanosoma cruzi*, responsible for Chagas disease, and *Leishmania mexicana*, which causes leishmaniasis, 3-(biphenyl-4-yl)-3-hydroxyquinuclidine acts as a chemotherapeutic agent by inhibiting the parasitic SQS to reduce endogenous sterol content and inhibit growth (Urbina et al., 2002). Unfortunately, 3-(biphenyl-4-yl)-3-hydroxyquinuclidine also inhibited mammalian SQS; however, in mammalian cell culture, this compound eradicated *T. cruzi* amastigotes, while the number of host cells remained unaffected. This was attributed to possible upregulation of the expression of LDL receptors to absorb cholesterol from the surrounding medium. Recently, studies on SQS from the filamentous fungus *Aspergillus flavus* have suggested this enzyme as a novel antifungal drug target (Song et al., 2019).

SQS comprises a catalytic N-terminal domain and a C-terminal domain, and the available crystal structures of protozoal, human and fungal SQSs share high overall homology (Liu et al., 2012; Malwal et al., 2022; Pandit et al., 2000). The C-terminal domain contains a transmembrane alpha-helix that anchors SQS to the membrane of the endoplasmic reticulum in vertebrates, yeasts and plants (Thompson et al., 1998), as well as to glycosomes and mitochondria in protozoal parasites (Urbina et al., 2002). In a study on *sqs* knock-out mutants of *Saccharomyces cerevisiae*, the activity of SQS could only be fully complemented with fungal *sqs* genes, or chimeric mammalian or plant *sqs* genes with a fungal-derived hinge region in the C-terminus (Linscott et al., 2016). Sequence comparisons revealed that this hinge region varies in a kingdom-specific manner; thus, selectively targeting the hinge region may be used to develop a fungal-specific inhibitor and warrants further investigation.

Currently, the structure of *A. flavus* SQS is the only fungal SQS structure available in the Protein Data Bank (PDB) (Malwal et al., 2022). *A. flavus* is most notorious for producing extremely carcinogenic aflatoxins (Klich, 2007), which contaminate food and feedstock, but can also cause invasive aspergillosis, especially in immune-compromised individuals (Rudramurthy et al., 2019). The World Health Organization has recently ranked the impact of fungal pathogens to guide research (World Health Organization, 2022), where *Candida albicans, Candida auris, Cryptococcus neoformans* and *Aspergillus fumigatus* form the critical priority pathogen group. In this study, we aim to produce high-quality SQS from *C. albicans* in *Escherichia coli* and purify the enzyme to homogeneity for future structure-based drug discovery studies.

## Materials and methods

### Amplification of sqs with PCR and cloning into expression vector

Genomic DNA was isolated from *C. albicans* SC5314 using the Quick-DNA Fungal/Bacterial Miniprep Kit (Zymo Research). The *sqs* gene was amplified from the genomic DNA with PCR using the KOD Hot Start DNA Polymerase kit (Merck), based on NCBI accession numbers XM_709367 and CP017624. The PCR reaction constituted 1X Buffer for KOD Hot Start DNA Polymerase, 1.5 mM MgSO_4_, dNTPs (0.2 mM each), sense primer (0.3 µM; 5’-<u>CATATG</u>GGCAAATTTTTACAATTATTATC-3’, *Nde*I recognition site underlined), anti-sense primers (0.3 µM; 5’-<u>GAATTC</u>AATAAAGGACAAAATA-AATTAAAC-3’, *Eco*RI recognition site underlined), 100 ng template DNA, and 0.3 U KOD Hot Start DNA Polymerase, to a total volume of 50 µl with deionised water. The thermocycling procedure was initiated with an initial polymerase activation step of 2 min at 95°C, and 40 cycles of denaturing (95°C, 20 s), annealing (50°C, 10 s), and extension (70°C, 40 s). A final elongation step at 70°C (5 min) was performed.

The PCR reaction was run on a 1% (w/v) agarose gel and the product corresponding to the expected size (1354 bp) was excised and purified using the GeneJET Gel Extraction kit (Thermo Scientific), according to the manufacturer’s instructions, with the exception of elution from the spin column using deionised water. The purified product was dried in a vacuum concentrator and resuspended in 13 µl deionised water. The PCR product was phosphorylated by adding 1 mM ATP, 1 U of T4 polynucleotide kinase (PNK, Thermo Scientific), 1X Buffer A, deionised water to a total volume of 20 µl, and incubated at 37°C for 20 min. The enzyme was deactivated by incubating at 80°C for 15 min. The phosphorylated product was ligated with pSMART HCKan (Lucigen) using 5 U T4 DNA ligase (Thermo Scientific), 1 μl 50% polyethylene glycol 4 000, 1X ligase buffer, 6 μl deactivated phosphorylation mixture, and 30 ng pSMART, to a total volume of 10 μl, and incubated for 1 h at room temperature. The ligation mixture was transformed into *E. coli* TOP10 cells (Thermo Scientific) using a heat-shock method and plated onto low-salt Luria Bertani (LB, 10 g.l^-1^ tryptone, 5 g.l^-1^ yeast extract, 5 g.l^-1^ NaCl) agar plates containing 30 mg.l^-1^ kanamycin. Plasmids were isolated using the GeneJET Plasmid Miniprep kit (Thermo Scientific) according to the manufacturer’s instructions, and screened with restriction enzyme digestion using *Eco*RI. Positive clones were verified with Sanger sequencing using the SL1 and SR2 primers. The *sqs* open reading frame was sub-cloned into pET-28b(+) (Novagen/Merck) using *Nde*I and *Eco*RI, which encodes an N-terminal 6xHis-tag. Plasmids were screened with restriction enzyme double-digestion, and verified with Sanger sequencing using the T7 promotor and terminator primers.

### Creation of N- and C-terminally truncated sqs variants

Based on a multiple-sequence alignment with crystallised variants of SQS from *T. cruzi* (PDB ID 3WCA) (Shang et al., 2014) and human SQS (PDB ID 1EZF, 3VJ8) (Liu et al., 2012; Pandit et al., 2000), two N-terminal truncations of SQS from *C. albicans* were designed, in which the first 21 residues were to be removed (Δ22) or the first 29 residues (Δ30). A homology model of SQS from *C. albicans* was generated using YASARA (Krieger & Vriend, 2014; Land & Humble, 2018), and C-terminal truncations were designed in which residues after the kingdom-specific hinge domain would be truncated to remove the α-helix membrane anchor. In ω384, residues 385-448 were to be removed, residues 391-448 for ω390, and residues 399-448 for ω398, respectively (Figure 2).

**Figure 2.**
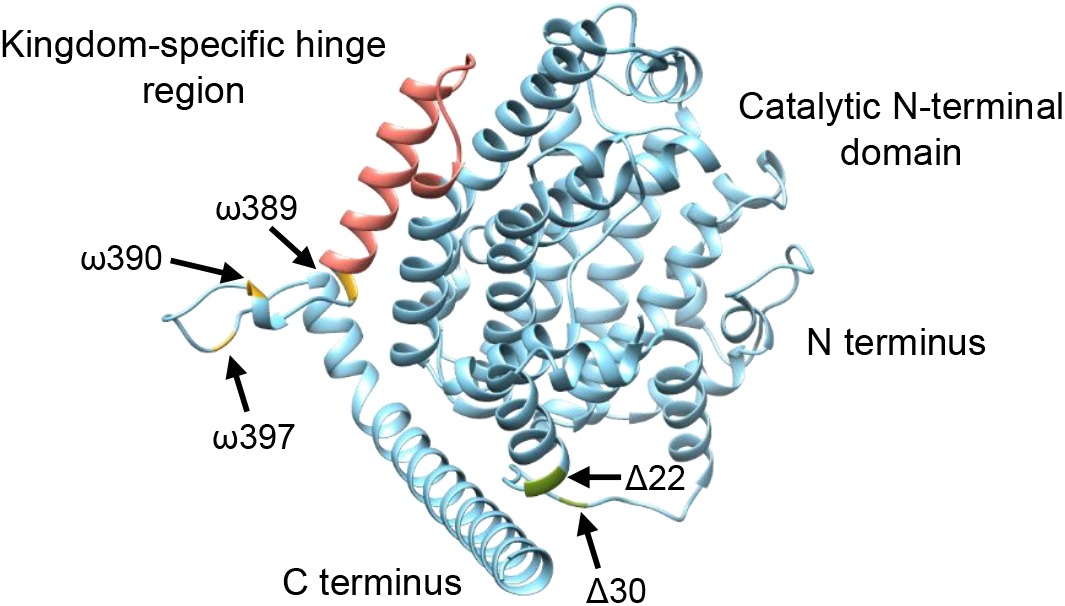
Homology model of *Candida albicans* squalene synthase (SQS) showing the kingdom-specific hinge region (salmon), C-terminal truncations (ω, gold) and N-terminal truncations (Δ, green).

N- or C-terminal regions were truncated using inverse PCR with pET-28b(+):*sqs* as template and primers that anneal on either side of the region to be removed (Table 1). PCRs were performed to remove either the N- or C-terminus using the KOD Hot Start DNA polymerase kit as previously described, with an annealing temperature of 57°C and an elongation time of 4 min. A *Dpn*I digestion (Thermo Scientific) was performed to remove the template DNA. The PCR mixtures were run on an agarose gel, and bands of ∼6600 bp were excised, purified, and dried in a vacuum concentrator. The linear PCR products were resuspended in 13 µl deionised water and circularised using a phosphorylation-ligation reaction comprising 15 U T4 PNK, 7.5 U T4 DNA ligase, 1 mM ATP and 1X T4 DNA ligase buffer to a total volume of 20 µl, which was incubated for 1 h at room temperature. The ligation mixture was transformed into *E. coli* TOP10 cells, plasmids were isolated and screened with restriction digestion, and successful truncation was verified with Sanger sequencing. N and C-terminal combinatorial truncated variants were created by using verified pET-28b(+):*sqs*Δ22 and pET-28b(+):*sqs*Δ30 as templates for C-terminal truncation PCRs.

**Table 1.**
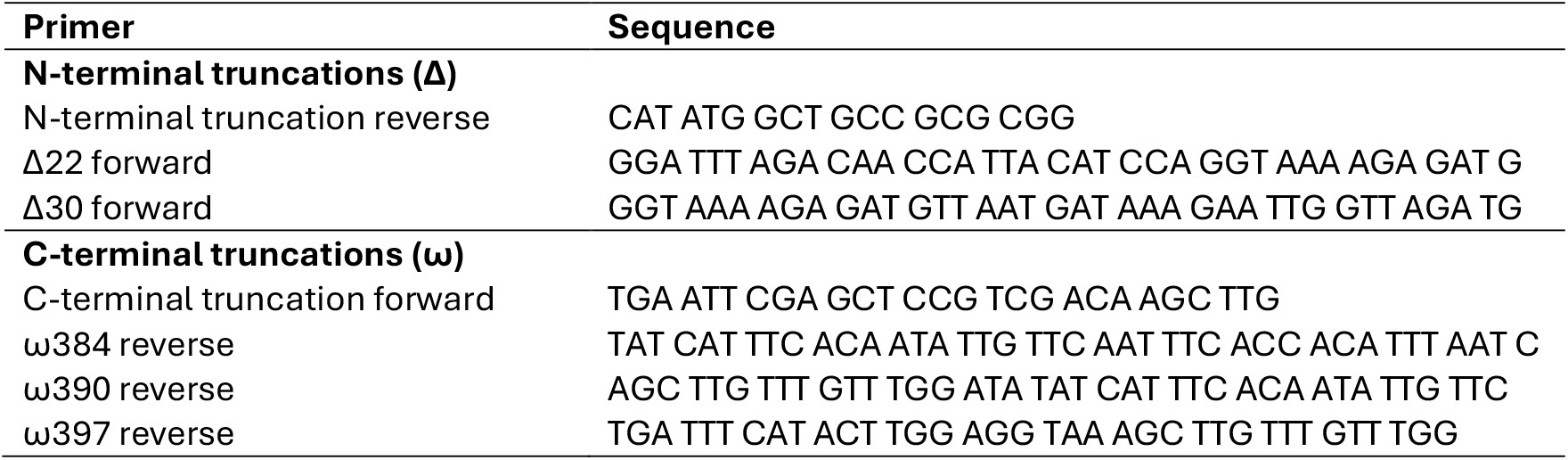
Primer sequences for the N- or C-terminal truncation of *sqs* in pET-28b(+).

### Protein expression and purification

The expression constructs were transformed into *E. coli* BL21 Gold-(DE3) (Agilent Technologies) for protein production. Colonies were collectively streaked and inoculated into 5 ml LB media (supplemented with kanamycin) to prepare a preculture by incubating at 37°C for 8 h with shaking. Four millilitres of the preculture were used to inoculate 200 ml of TB auto-induction media (12 g.l^-1^ tryptone, 24 g.l^-1^ yeast extract, 0.8% (v/v) glycerol, 5 g.l^-1^ lactose, 0.15 g.l^-1^ glucose, 17 mM KH_2_PO_4_, 73 mM K_2_HPO_4_, 3.75 g.l^-1^ aspartate, 1.2 g.l^-1^ NaOH, 2 mM MgSO_4_) in a 1 l Erlenmeyer flask, which was incubated at 20°C for 40 h. Cells were harvested by centrifugation at 7000 x*g* and 4°C for 10 min. The cells were resuspended in immobilised metal-affinity chromatography (IMAC) binding buffer, containing 50 mM Tris-HCl pH 7.4, 30 mM imidazole, and 500 mM NaCl, in a 1 g cells:5 ml buffer ratio. Cells were disrupted with a One Shot Cell Disruptor (Constant Systems) at 30 kpsi and the lysate was centrifuged at 20 000 x*g* and 4°C for 20 min to remove the insoluble debris. The supernatant was further ultracentrifuged at 90 000 x*g* and 4°C for 1 h to remove the membranes.

The soluble fractions were loaded onto His GraviTrap columns (Cytiva), equilibrated with binding buffer, for IMAC batch-purification. The columns were washed with 2 x 25 ml binding buffer, and the bound proteins were eluted with 2.5 ml elution buffer (50 mM Tris pH 7.4, 500 mM imidazole, and 500 mM NaCl). Protein expression and purification were evaluated with SDS-PAGE.

Expression of SQS variants ω397 and Δ30ω397 were upscaled for purification to produce 3 l of culture each. Cells were lysed with a Continuous Flow Cell Disruptor (Constant Systems) at 30 kpsi and 4°C. The proteins were purified using HisTrapFF columns (5 ml, Cytiva) on an ÄKTA Pure chromatography system (Cytiva). The soluble fractions were loaded on equilibrated columns, and unbound proteins were removed with 20 column volumes of binding buffer. The proteins were eluted with a linear increasing gradient of imidazole (30-500 mM) over 20 column volumes. The elution of proteins was monitored spectrophotometrically by absorption at 280 nm. Fractions of interest were pooled and concentrated with a 30 kDa MWCO Amicon centrifugal filter concentrator (Merck) at 3 500 x*g* and 4°C. Size-exclusion chromatography was performed with a Sephacryl HR100 column (Cytiva) using buffer comprising 50 mM Tris pH 7.4 and 500 mM NaCl. Protein concentration was determined with the Pierce BCA Protein Assay kit (Thermo Scientific) using bovine serum albumin as standard.

### Crystallisation trials

Sitting drop vapour diffusion crystallisation trials were set up in 96-well plates with the Oryx Nano crystallisation robot (Douglas Instruments) using the Crystal 1&2, PEGRx 1&2, PEG/Ion 1&2, and SaltRx 1&2 screens (Hampton Research). SQS variants ω397 and Δ30ω397 were used at a concentration of 4 - 8 mg.ml^-1^ with a 1:1 ratio of protein:reservoir solution to produce drops of 1 µl in 3-well SWISSCI crystallisation plates. Crystallisation plates were incubated at 16°C.

## Results

### Cloning and truncation library construction

Based on previous studies on the heterologous production and crystallisation of human (Liu et al., 2012; Thompson et al., 1998) and trypanosomal (Shang et al., 2014) SQS, we anticipated that successful heterologous production, purification and crystallisation of *C. albicans* SQS would require expression of truncated *sqs* variants. The full-length *sqs* open reading frame was successfully amplified from the genomic DNA of *C. albicans* by PCR, cloned into the pSMART cloning vector, and sub-cloned into pET-28b(+). By using pET-28b(+):*sqs* as a template, two N-terminal truncation variants (Δ22 and Δ30), three C-terminal truncation variants (ω384, ω390, ω397), and six combinatorial variants, with regions truncated on both N- and C-termini (Δ22ω384, Δ22ω390, Δ22ω397; Δ30ω384, Δ30ω390, Δ30ω397) were constructed.

### Test expression and IMAC batch-purification

Test expression of the library of *sqs* variants was performed in *E. coli* using TB auto-induction media. No expression of the full-length *sqs* or N-terminally truncated variants was observed. This may be due to inclusion of the C-terminal membrane anchor in the variants. All C-terminally and combinatorially truncated variants were successfully expressed, producing proteins corresponding approximately to the expected molecular weights (42.0-45.7 kDa) (Figure 3). Expression levels were similar between the variants.

**Figure 3.**
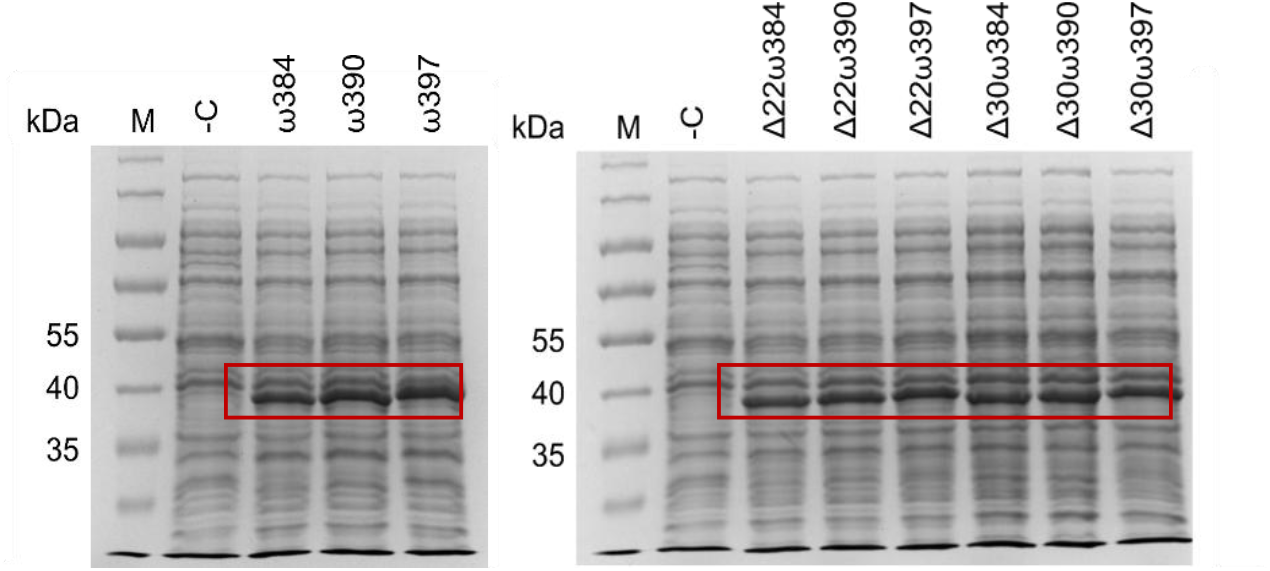
SDS-PAGE analysis of the expression of SQS variants in *E. coli*. ‘ω’ denotes C-terminal truncations and ‘Δ’ denotes N-terminal truncations. ‘-C’, negative control (empty pET-28b(+)); ‘M’, PageRuler Prestained Protein Ladder. Expected sizes for SQS variants range from 42.0-45.7 kDa.

All expressed variants were batch-purified using His GraviTrap columns, and bound to and eluted from the columns with high yield (Figure 4). However, several contaminating proteins were still present.

**Figure 4.**
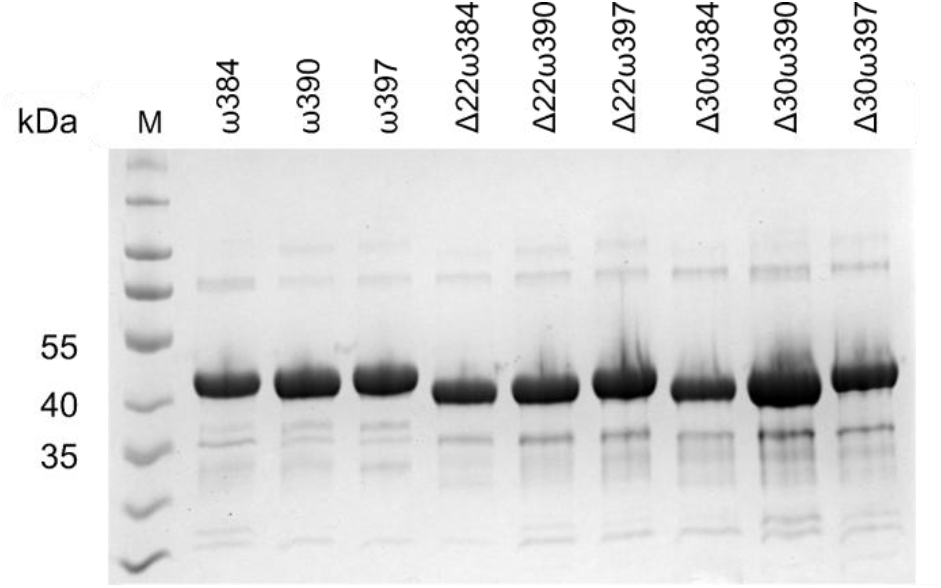
SDS-PAGE analysis of SQS variants batch-purified using IMAC. ‘ω’ denotes C-terminal truncations and ‘Δ’ denotes N-terminal truncations. Expected sizes for SQS variants range from 42.0-45.7 kDa.

### Large-scale expression and purification by immobilised metal-affinity (IMAC) and size-exclusion chromatography (SEC)

Two SQS variants were selected for upscaling and purification, ω397 and Δ30ω397. These represented the largest C-terminally truncated SQS variant versus its most truncated N-terminal counterpart. The proteins remained soluble after ultracentrifugation, which confirmed successful separation from the membranes, and the yield without the addition of detergent to the binding buffer was similar to the yield in the presence of Triton-X100, Tween 20 or CHAPS (1% (w/v)). Elution from the HisTrapFF column using an imidazole gradient significantly improved the purity of the protein, which was followed by SEC to ensure homogeneity (Figure 5).

**Figure 5.**
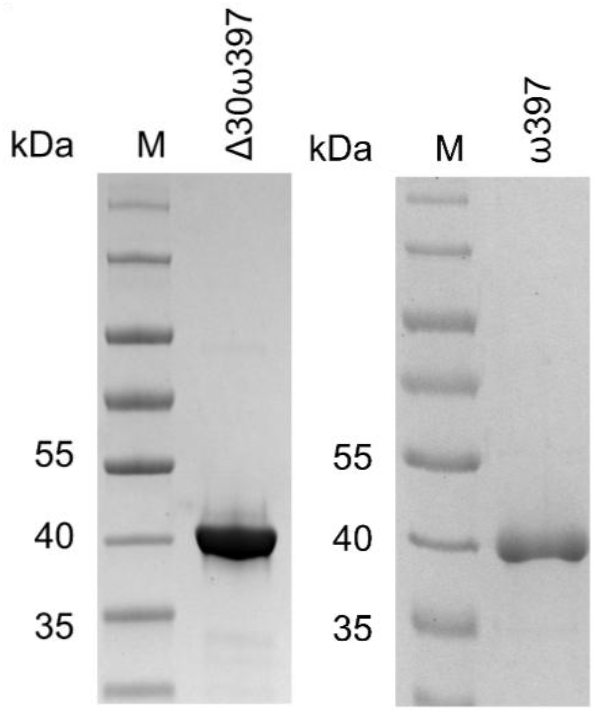
SDS-PAGE analysis of the purification of SQS variants Δ30ω397 and ω397 by size-exclusion chromatography.

SEC elution profiles indicated sample heterogeneity with possible aggregation for both variants; however, this was more pronounced for ω397. A peak corresponding to the monomer was observed for Δ30ω397, and fractions could be selectively pooled, but a defined peak for the ω397 monomer could not be observed. The SQS variants were very sensitive towards NaCl concentration, with immediate precipitation when buffer with a concentration of 10 – 250 mM was used. Furthermore, cell lysis, centrifugation, IMAC, and SEC were performed on the same day as precipitation was observed when the protein was stored at 4°C in IMAC elution buffer overnight.

Crystallisation trials using the 96-well sitting drop vapour diffusion method are currently underway, but no crystals have been produced. Generally, conditions with ω397 tended to precipitate soon after crystallisation despite lowering the protein concentration, while Δ30ω397 remained soluble or produced amorphous precipitate.

## Discussion and conclusion

*Candida albicans* is a pathogenic yeast that can cause invasive candidiasis with mortality rates of 20-50% despite antifungal treatment (World Health Organization, 2022). A limited number of antifungal treatment regimens are available clinically, some of which are difficult to administer (e.g., only by IV), have severe toxicity, or are not readily available in all countries. Moreover, resistance towards these compounds by pathogenic fungi has emerged globally and is steadily increasing (Lopes & Lionakis, 2022). In South Africa, infectious fungal disease significantly contributes to the public health burden due to the high HIV/AIDS incidence, and drug-resistant *Candida* species have been isolated from patients for over a decade (Dos Santos Abrantes et al., 2014).

Several avenues for antifungal drug discovery are being investigated, but structure-based techniques have been under-utilised in the search for novel antifungal medicines. The divergence in sterol composition and biosynthesis between fungal and mammalian cells poses an ideal target for drug discovery efforts. However, many enzymatic steps are catalysed by homologous enzymes in the two kingdoms, and cross-inhibition may possibly be observed, as seen in the study by Urbina and co-workers (2002). Therefore, targeting fungal-specific enzymes, or fungal-specific features on highly homologous enzymes, may be advantageous.

Squalene synthase (SQS) catalyses a key reaction in the synthesis of ergosterol and has been identified as a possible target for antifungal drug development (Song et al., 2019). A hinge region in the C-terminus has been shown to vary in a kingdom-specific manner (Linscott et al., 2016), and can possibly be exploited to develop non-active site antifungal inhibitors despite the high overall homology between the human and fungal SQSs.

Here, we heterologously produced and purified SQS from *C. albicans* for future structural studies. Nine SQS variants with either C-terminal or N&C-terminal truncations were produced in *E. coli* and purified with IMAC. Two variants were purified to near-homogeneity with IMAC and SEC, but showed possible aggregation; therefore, future studies will focus on buffer optimisation to improve monodispersity. Vapour-diffusion crystallisation trials are currently underway with the apo form of *C. albicans* SQS, and co-crystallisation with ligands such as S-*thiolo*-farnesyl pyrophosphate can be explored.

## Acknowledgements

This work is based on the research supported in part by the National Research Foundation of South Africa (TTK200401511013) and the University of the Free State, South Africa. The authors would like to acknowledge the experimental contribution of Miss Siyavuya Sidla and Mr Schalk Campher to this work.

